# Anticancer efficacy of KRASG12C inhibitors is potentiated by PAK4 inhibitor KPT9274 in preclinical models of KRASG12C mutant pancreatic and lung cancers

**DOI:** 10.1101/2023.03.27.534309

**Authors:** Husain Yar Khan, Misako Nagasaka, Amro Aboukameel, Osama Alkhalili, Md. Hafiz Uddin, Sahar Bannoura, Yousef Mzannar, Ibrahim Azar, Eliza Beal, Miguel Tobon, Steve Kim, Rafic Beydoun, Erkan Baloglu, William Senapedis, Bassel El-Rayes, Philip A. Philip, Ramzi M. Mohammad, Anthony F. Shields, Mohammed Najeeb Al-Hallak, Asfar S. Azmi

**Affiliations:** Barbara Ann Karmanos Cancer Institute, Department of Oncology, Wayne State University School of Medicine, Detroit MI 48201, USA; University of California Irvine School of Medicine, Orange CA 92868, USA; Chao Family Comprehensive Cancer Center, Orange, CA 92868, USA; Division of Neurology, Department of Internal Medicine, St. Marianna University, Kawasaki, Japan; Karyopharm Therapeutics, Newton MA 02459, USA; University of Alabama, Birmingham, Al USA; Henry Ford Health, Detroit, MI 48201

**Keywords:** KRASG12C, PAK4, KPT9274, sotorasib, adagrasib, combination therapy

## Abstract

KRASG12C inhibitors have revolutionized the treatment landscape for cancer patients harboring the G12C mutant isoform of KRAS. With the recent FDA approval of sotorasib and adagrasib, patients now have access to more promising treatment options. However, patients who receive these agents as a monotherapy usually develop drug resistance. Thus, there is a need to develop logical combination strategies that can delay or prevent the onset of resistance and simultaneously enhance the antitumor effectiveness of the treatment regimen. In this study, we aimed at pharmacologically targeting PAK4 by KPT9274 in combination with KRASG12C inhibitors in KRASG12C mutant pancreatic ductal adenocarcinoma (PDAC) and non–small cell lung cancer (NSCLC) preclinical models. PAK4 is a hub molecule that links several major signaling pathways and is known for its tumorigenic role in mutant Ras-driven cancers. We assessed the cytotoxicity of PAK4 and KRASG12C inhibitors combination in KRASG12C mutant 2D and 3D cellular models. KPT9274 synergized with both sotorasib and adagrasib in inhibiting the growth of KRASG12C mutant cancer cells. The combination was able to reduce the clonogenic potential of KRASG12C mutant PDAC cells. We also evaluated the antitumor activity of the combination in a KRASG12C mutant PDAC cell line-derived xenograft (CDX) model. Oral administration of a sub-optimal dose of KPT9274 in combination with sotorasib (at one-fourth of MTD) demonstrated significant inhibition of the tumor burden (*p* = 0.002). Similarly, potent antitumor efficacy was observed in an NSCLC CDX model where KPT9274, acting as an adjuvant, prevented tumor relapse following the discontinuation of sotorasib treatment (*p* = 0.0001). KPT9274 and sotorasib combination also resulted in enhanced survival. This is the first study showing that KRASG12C inhibitors can synergize with PAK4 inhibitor KPT9274 both *in vitro* and *in vivo* resulting in remarkably enhanced antitumor activity and survival outcomes.

**Significance:** KRASG12C inhibitors demonstrate limited durable response in patients with KRASG12C mutations. In this study, combining PAK4 inhibitor KPT9274 with KRASG12C inhibitors has resulted in potent antitumor effects in preclinical cancer models of PDAC and NSCLC. Our results bring forward a novel combination therapy for cancer patients that do not respond or develop resistance to KRASG12C inhibitor treatment.

## INTRODUCTION

KRAS is one of the most common oncogenes in human cancers, with a particularly high mutation rate in pancreatic, colorectal, and lung cancers [1]. Given its high prevalence in various malignancies, extensive efforts have been dedicated to exploring the potential for direct inhibition of mutated KRAS. A number of small molecule inhibitors, including sotorasib (AMG510) and adagrasib (MRTX849), have been developed that target the KRASG12C mutant protein in cancer cells [2,3]. These inhibitors rely on the mutant cysteine to bind irreversibly to the switch II pocket on the GDP-bound KRAS protein, resulting in the trapping of KRAS in the inactive state and the inhibition of KRAS-dependent signaling [4].

Clinical trials are currently underway to assess the effects of these covalent inhibitors of KRASG12C on cancer patients. Preliminary trials of sotorasib and adagrasib have reported encouraging results [5,6]. The emergence of these new covalent inhibitors has presented an unparalleled opportunity to target the critical KRAS mutation. Notably, in 2021, the FDA granted accelerated approval of sotorasib for treating patients with KRASG12C mutated locally advanced or metastatic non-small cell lung cancer [7]. Recently, adagrasib became the second KRAS inhibitor to receive an FDA approval [8].

While studies on KRASG12C inhibitors have demonstrated encouraging outcomes, the fact that some patients do not respond to these therapies, show limited durable response and experience recurrence suggests that the issue of intrinsic or acquired resistance is an inevitable challenge for such molecular targeted therapies. This resistance is likely due to the activation of an alternative pathway or the selection of minor drug-resistant mutants. Therefore, circumventing drug resistance by developing novel combination therapies that can maximize the therapeutic potential of these KRASG12C drugs is the need of the hour.

A relatively less explored effector of RAS oncogenic signaling is PAK4 which belongs to a group of serine/threonine kinases called p21-activated kinases (PAK). PAK4 has emerged as an appealing target for cancer therapy due to its strategic role as a hub molecule linking major signaling pathways, including the KRAS-MAPK pathway [9, 10]. PAK4 is often upregulated in various types of cancer, including NSCLC, prostate, gastric and breast cancers [11-14]. In cancers such as pancreatic, ovarian and oral squamous cell carcinoma, amplification of the PAK4 gene is frequently observed [15-17]. PAK4 plays a critical role in controlling cell proliferation, survival, invasion, metastasis, epithelial-mesenchymal transition, and drug resistance both *in vitro* and *in vivo*, ultimately contributing to cancer progression [10-12, 18-20]. It is noteworthy that PAK4, as opposed to other PAK isoforms, can transform normal cells [19]. A recent study demonstrated that PAKs were activated in KRASG12C inhibitor-resistant cancer cells and PAKs mediate this resistance through the activation of MAPK and PI3K pathways [21]. PAK4 signaling has also been reported to play a critical role in resistance to chemotherapy in gastric and cervical cancers [22,23]. Moreover, targeting PAK4 has been shown to inhibit Ras-mediated oncogenic signaling [24].

KPT9274, a small molecule inhibitor that targets PAK4, has been studied as a potential cancer therapy in preclinical and early-stage clinical trials [20]. In this study, we report for the first time that KRASG12C inhibitor-resistant cancer cells show sensitivity toward KPT9274, providing a rationale for testing PAK4 inhibitor in combination with KRASG12C inhibitors as an effective combination therapy. Using KRASG12C mutant preclinical cancer models, we demonstrate that KPT9274 synergizes with KRASG12C inhibitors and enhances their anticancer activity when used in combination as well as an adjuvant.

## MATERIALS AND METHODS

### Cell lines, drugs, and reagents

MIA PaCa-2, NCI-H358 were purchased from American Type Culture Collection (ATCC, Manassas, VA, USA) in 2012 and 2021, respectively. NCI “Rasless” mouse embryonic fibroblast (MEF) cell lines (KRAS 4B G12C and KRAS 4B G12D) were obtained from the National Cancer Institute (Rockville, MD, USA) in 2019. MIA PaCa-2 and the MEF cell lines were maintained in DMEM (Thermo Fisher Scientific, Waltham, MA, USA), while NCI-H358 was maintained in RPMI1640 (Thermo Fisher Scientific, Waltham, MA, USA), supplemented with 10% fetal bovine serum (FBS), 100 U/mL penicillin, and 100 μg/mL streptomycin in a 5% CO_2_ atmosphere at 37 °C. All experiments were performed within 20 passages of the cell lines. AMG510, MRTX849 (Selleck Chemical LLC, Houston, TX) and KPT9274 (Karyopharm Therapeutics, Newton, MA, USA) were dissolved in DMSO to make 10 mM stock solutions. The drug control used for *in vitro* inhibitor experiments was cell culture media containing 0.1% DMSO.

### Cell viability assay and synergy analysis

Cells were seeded in 96-well culture plates at a density of 3 × 10^3^ cells per well. The growth medium was removed after overnight incubation and replaced with 100 µL of fresh medium containing the drug at various concentrations serially diluted from stock solution using OT-2 liquid handling robot (Opentrons, Queens, NY, USA). After 72 hours of exposure to the drug, MTT (3-(4,5-dimethylthiazol-2-yl)-2,5-diphenyltetrazolium bromide) assay was performed according to the procedure described previously [25]. Using the cell proliferation data (six replicates for each dose), IC_50_ values were calculated using the GraphPad Prism 4 software. For the synergy analysis, cells were treated with three different concentrations of either MRTX849/AMG510, or KPT9274, or a combination of KPT9274 with MRTX849/AMG510 at the corresponding doses for 72 hours (six replicates for each treatment). The ratio of drug concentration was kept constant across all the three dose combinations tested. Cell growth index was determined using MTT assay. The resulting cell growth data was used to calculate combination index (CI) values by the CalcuSyn software (Biosoft, Cambridge, UK).

### Colony formation assay

MIA PaCa-2 cells were seeded at a density of 500 cells per well in six well plates and exposed to single agent or combination drug treatments for 72 h. At the end of the treatment, drug containing media was removed and replaced with fresh media. The plates were incubated in the CO_2_ incubator for an additional ten days. After the incubation was over, media was removed from the wells of the plates and the colonies were fixed with methanol and stained with crystal violet for 15 minutes. The plates were then washed and dried before colonies were photographed.

### Spheroid formation and disintegration assays

MIA PaCa-2 and NCI-H358 cells were trypsinized, collected as single cell suspensions using cell strainer and resuspended in 3D Tumorsphere Medium XF (PromoCell, Heidelberg, Germany). For tumor formation assay, 1,000 cells were plated in each well of ultra-low attachment 6-well plate (Corning, Durham, NC, USA) and treated with either KPT9274, or MRTX849, or a combination of KPT9274 with MRTX849 twice a week for one week. For the spheroid disintegration assay, 200 cells were plated in each well of an ultra-low attachment round bottom plate (Corning, Durham, NC, USA). Media was replenished every 3 days and spheroid growth was monitored. Once sizable spheroids formed, they were subjected to treatment with the drugs for three days. At the end of the treatment, spheroids were photographed under an inverted microscope.

### KRASG12C mutant cell-derived tumor xenograft (CDX) studies

*In vivo* studies were conducted under Wayne State University’s Institutional Animal Care and Use Committee (IACUC) approved protocol in accordance with the approved guidelines. Experiments were approved by the institute’s IACUC (#18-12-0887 for PDAC *in vivo* study and #22-01-4355 for NSCLC *in vivo* study).

### PDAC (MIA PaCa-2) CDX model

Post adaptation in our animal housing facility, 4-5 weeks old female ICR-SCID mice (Taconic Biosciences, Rensselaer, NY) were subcutaneously implanted with MIA PaCa-2 cells. 1 × 10^6^ cells suspended in 200 μL PBS were injected unilaterally into the left flank of donor mice using a BD 26Gx 5/8 1ml Sub-Q syringe. Once the tumors reached about 5-10% of the donor mice body weight, the donor mice were euthanized, tumors were harvested, and fragments were subsequently implanted into recipient mice. Seven days post transplantation, the recipient mice were randomly divided into four groups of 6 mice each and received either vehicle, or KPT9274 (100 mg/Kg QD × 5 × 3 weeks), or sotorasib (25 mg/Kg QD × 5 × 3 weeks), or their combination by oral gavage. Tumor volumes were determined using the formula 0.5 _×_ *L* _×_ *W*^2^ in which *L* refers to length and *W* refers to width of each tumor. On completion of drug dosing, tumor tissue from control or treatment groups were used for IHC and molecular analysis.

### NSCLC (NCI-H358) CDX model

NCI-H358 cells were washed in PBS, then suspended in cold PBS at a concentration of 100,000 cells per 100 μL. The cell suspensions were mixed with an equal volume of Cultrex extracellular matrix (ECM; Trevigen; #3432-005-01) and kept on ice. Female ICR-SCID mice, aged 4-5 weeks, were injected with the mixture of ECM and cells into the left flank of each mouse. For the efficacy study, mice were randomized into four groups of 5 mice each. Mice were orally administered sotorasib (25 mg/Kg QD x 5) for 15 days in the single agent and combination groups, while KPT9274 (150 mg/Kg QD x 5) was dosed for 26 days in both the groups. For the survival study, mice unilaterally transplanted with NCI-H358 cells were randomly divided into four treatment groups of 7 mice each. Sotorasib and KPT9274 were administered at the abovementioned doses for three and five weeks, respectively, by oral gavage.

### Immunostaining

Paraffin sections of the MIA PaCa-2 derived tumors were processed and stained with H&E and antibodies in the core facility at the Department of Oncology, Wayne State University/Karmanos Cancer Institute. The following antibodies were used for immunohistochemistry staining: anti-Ki67 and anti-KRAS at 1:100 dilution, and anti-PAK4 at 1:50 dilution.

### Preparation of total protein lysates and Western Blot analysis

For total protein extraction, tumor tissues were lysed in RIPA buffer and protein concentrations were measured using BCA protein assay (PIERCE, Rockford, IL, USA). A total of 40 μg protein lysate from treated and untreated cells was resolved on 10% SDS-PAGE and transferred onto nitrocellulose membranes. The membranes were incubated with the following primary antibodies (Cell Signaling Technology, Danvers, MA, USA) at 1:1000 dilution in 3% non-fat dry milk: anti-phospho-Erk ½ (# 4370), anti-Erk ½ (# 9102), anti-DUSP6 (# 50945), anti-CDK4 (# 12790), anti-CDK6 (# 13331), anti-cyclin D1 (# 55506). While anti-GAPDH (# sc-47724; Santa Cruz Biotechnology, Santa Cruz, CA, USA) was used at a dilution of 1:3000. Incubation with 1:2000 diluted HRP-linked secondary antibodies (# 7074/7076; Cell Signaling, Danvers, MA, USA) in 3% non-fat dry milk was subsequently performed at room temperature for 1 hour. The signal was detected using the ECL chemiluminescence detection system (Thermo Fisher Scientific, Waltham, MA, USA).

### RNA isolation and mRNA real-time RT-qPCR

Total RNAs from mouse tumors were extracted and purified using the RNeasy Mini Kit and RNase-free DNase Set (QIAGEN, Valencia, CA) following the protocol provided by the manufacturer. The expression levels of PAK4 in the mouse tumor tissues were analyzed by real-time RT-qPCR using High-Capacity cDNA Reverse Transcription Kit and SYBR Green Master Mixture from Applied Bio-systems (Waltham, MA, USA). The conditions and procedure for RT-qPCR have been described previously [29]. Sequences of primers used are as below: Pak4_F - GTGCAAGAGAGCTGAGGGAG Pak4_R - ATGCTGGTGGGACAGAAGTG

### Statistical analysis

The student *t* test was used to compare statistically significant differences. Wherever suitable, the experiments were performed at least three times. The data were also subjected to unpaired two-tailed Student *t* test wherever appropriate and *P < 0*.*05* was considered statistically significant.

### Data Availability

The data generated in this study are available upon request from the corresponding author.

## RESULTS

### KPT9274 induces growth inhibition in PDAC cells resistant to KRASG12C inhibitor

KRASG12C inhibitor-resistant cell lines were generated in our lab as described previously [26]. To establish the durability of drug-resistance, we compared the IC_50_ values of sotorasib-resistant cell line (MIA-AMG-R) that we developed earlier with the sotorasib-sensitive parental cell line (MIA PaCa-2). We observed more than 42-fold increase in the IC_50_ of sotorasib for the MIA-AMG-R cells, confirming that these cells still maintain their resistance to sotorasib even in the absence of sotorasib from their growth medium (**Figure 1A**). Subsequently, this drug-resistant cell line was treated with KPT9274 and was found to be sensitive to KPT9274-induced cell growth inhibition (**Figure 1B**). This establishes that the KRASG12C inhibitor-resistant cancer cells can potentially respond to the PAK4 inhibitor KPT9274.

**Figure 1:**
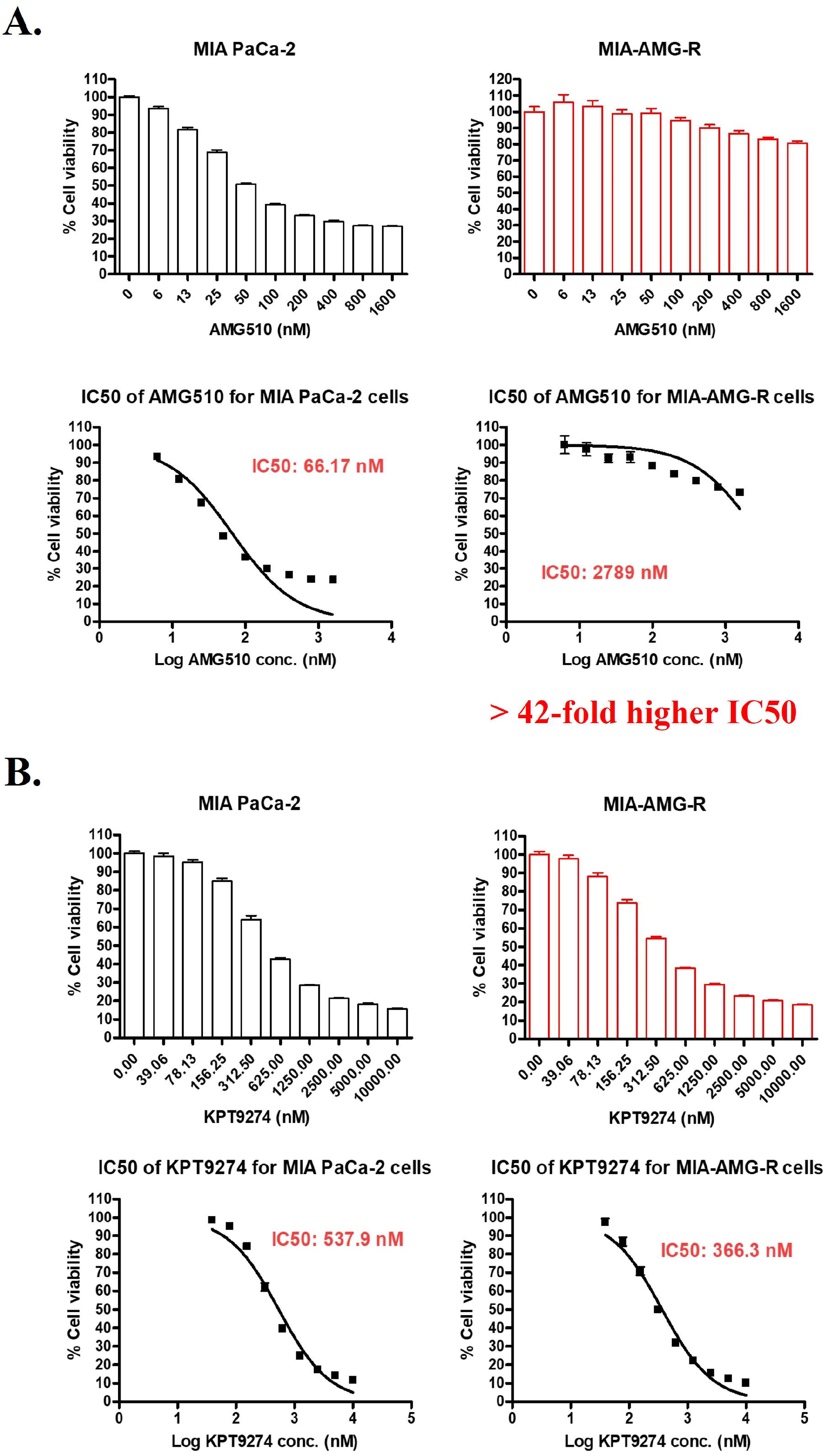
KPT9274 induces growth inhibition in KRASG12C inhibitor resistant cancer cells. (**A**) KRASG12C mutant MIA PaCa-2 cells exposed to incremental doses of AMG510 in long term cell culture eventually developed drug-resistance (MIA-AMG-R) as shown by their unresponsiveness to drug treatment in MTT assay and > 42-fold increase in the drug IC_50_ value compared to parental MIA PaCa-2 cells. (**B**) AMG510-resistant MIA PaCa-2 cell line (MIA-AMG-R) show sensitivity toward KPT9274 induced growth inhibition. Parental as well as resistant cells were treated with KPT9274 for 72h and MTT assay was performed as described in Methods. All results are expressed as percentage of control ± S.E.M of six replicates.

Next, we used Synthetic Lethal analysis via Network topology (SLant) to predict human synthetic lethal (SSL) interactions [27]. This approach identifies and leverages conserved patterns in the topology of protein interaction networks for its predictions. Using this approach, we have obtained the experimentally validated synthetic lethal interactions of PAK4 from the Slorth database (http://slorth.biochem.sussex.ac.uk/welcome/index) and found the interaction of PAK4 with KRAS to be synthetic lethal (**Supplementary Figure 1**).

### Combining KPT9274 with KRASG12C inhibitors synergistically suppresses the proliferation of KRAS G12C mutant cells

KRASG12C mutant MIA PaCa-2 (PDAC) and NCI-H358 (NSCLC) cells were exposed to MRTX849/AMG510 and KPT9274 at different dose combinations. As shown in **Figure 2A-B**, all three dose combinations tested demonstrated synergistic inhibition of MIA PaCa-2 cell proliferation (CI value < 1). Similar synergistic effects (CI < 1) in suppressing cell growth were also observed with the NCI-H358 cells treated with different dose combinations of the two drugs (**Figure 2C-D**). This drug combination was also tested on NCI “Rasless” MEFs carrying KRASG12C or G12D mutations. KPT9274 synergized with MRTX849 at all dose combinations yielding suppressed growth of KRASG12C mutant MEFs (**Supplementary Figure 2A**). As expected, the KRAS G12D MEFs (KRAS4B G12D) were refractory to any such growth inhibition (**Supplementary Figure 2B**).

**Figure 2:**
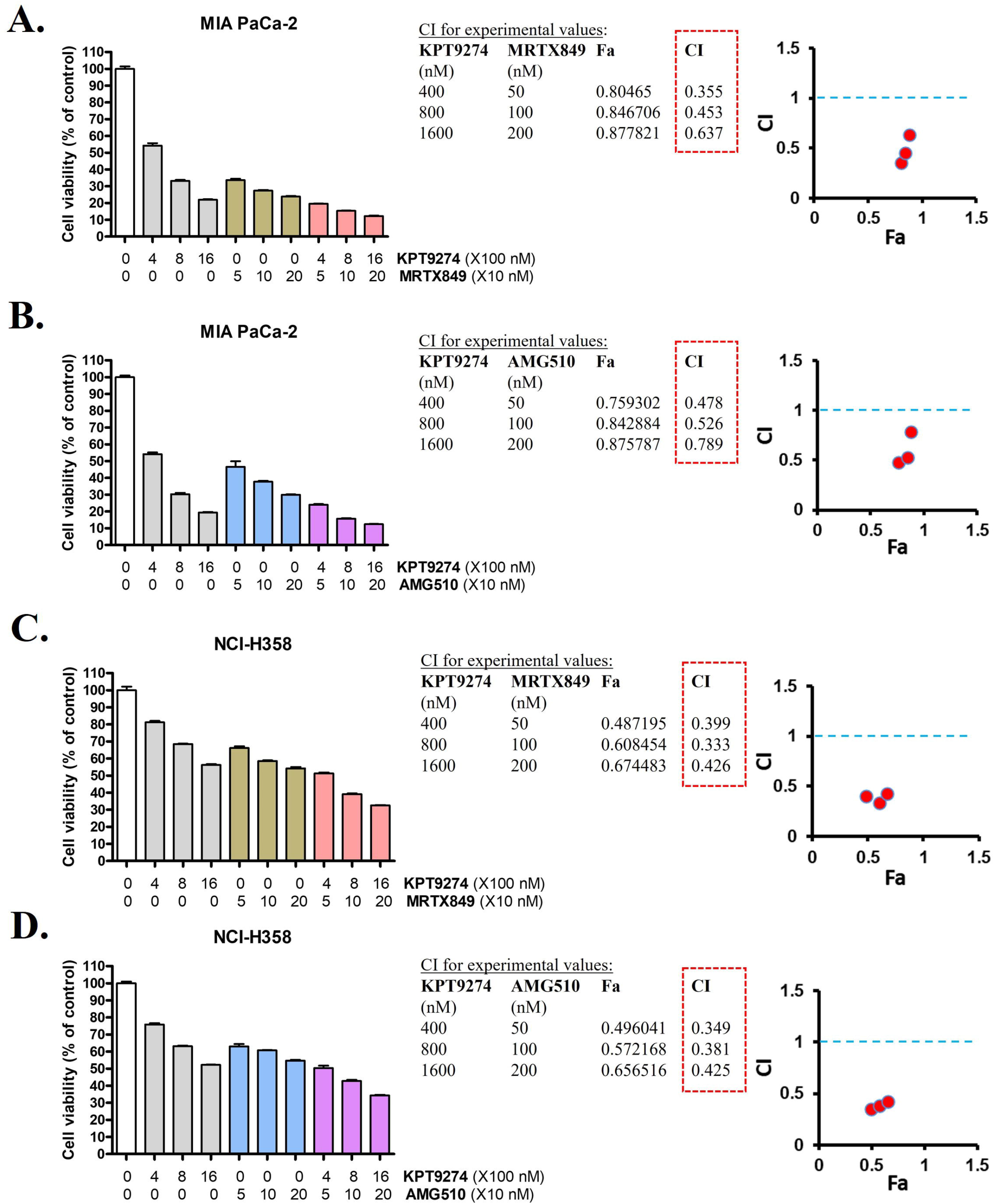
KPT9274 and KRASG12C inhibitors show synergistic effects on the inhibition of cell proliferation *in vitro*. Different dose combinations of KPT9274 with MRTX849 (**A, C**) and AMG510 (**C, D**) synergistically inhibit the proliferation of KRASG12C mutant PDAC cell line MIA PaCa-2 and NSCLC cell line NCI-H358, as indicated by combination index (CI) values less than 1. Cell viability was determined by MTT assay and synergy analysis was performed by using Calcusyn software. All results are expressed as percentage of control ± S.E.M of six replicates.

### Combination of KPT9274 with KRASG12C inhibitor reduces the clonogenic potential and effectively disrupts spheroids

The combinations of KPT9274 with MRTX849 were evaluated for their effects on the colony formation ability of MIA PaCa-2 cells. Results of a clonogenic assay clearly demonstrate that the combination treatments of KPT9274 with the KRASG12C inhibitor resulted in a decrease in colony numbers as well as the size of colonies formed by MIA PaCa-2 cells (**Figure 3A**). These findings further underscore the efficacy of this combination approach in targeting KRASG12C mutant cancer cells *in vitro*.

**Figure 3:**
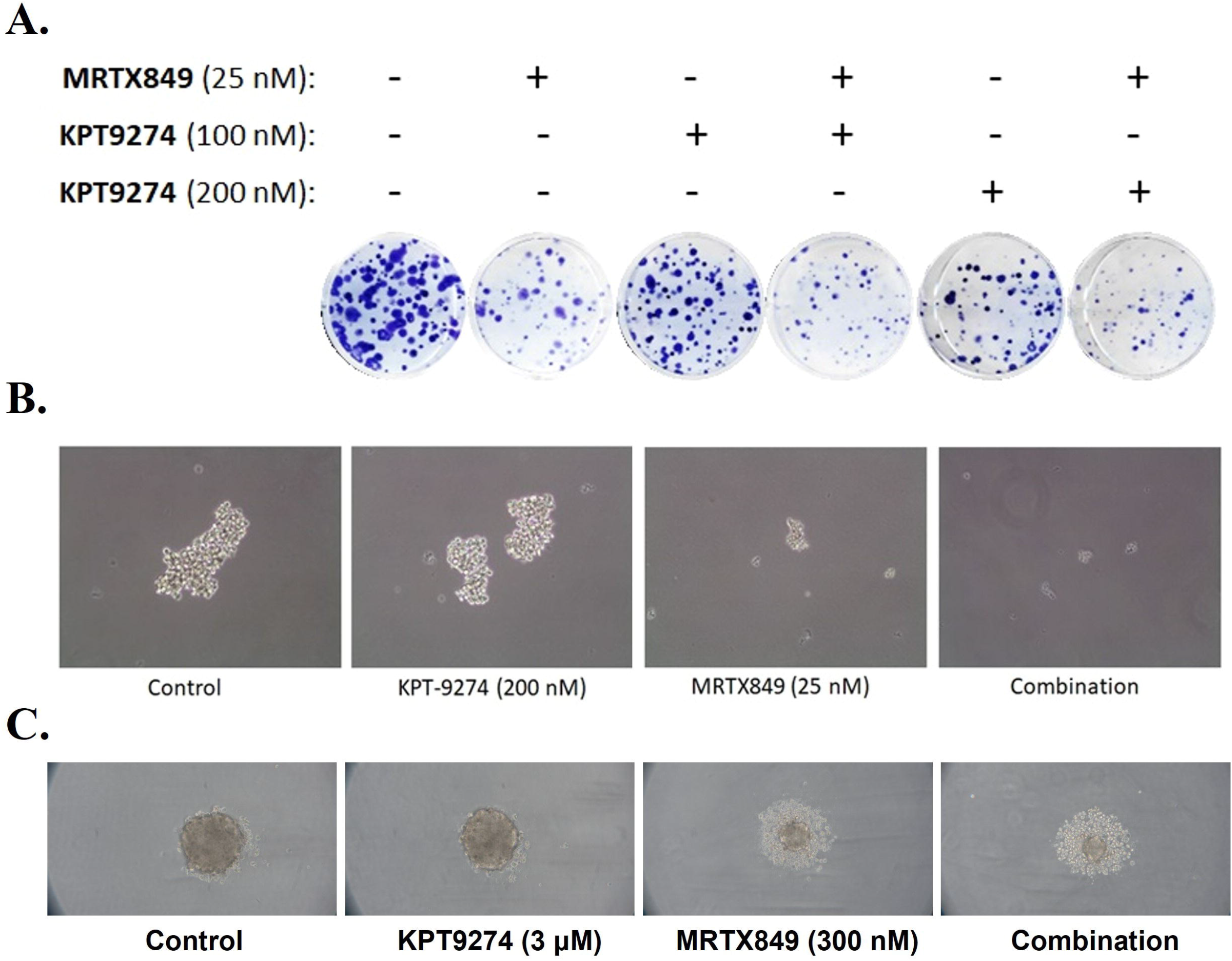
KPT9274 and KRASG12C inhibitor combination reduces the clonogenic potential, prevents spheroid formation and induces spheroid disintegration. (**A**) Combination of KPT9274 with MRTX849 inhibits the ability of KRASG12C mutant PDAC cells to form colonies. MIA PaCa-2 cells seeded in 6 well plates (500 cells/well) were treated with indicated doses of the drugs for 72 hours and the colonies were fixed and stained after one week. KPT9274 and MRTX849 combination treatment (**B**) suppresses spheroid formation in 3D cultures of MIA PaCa-2 cells and (**C**) enhances disintegration of spheroids formed by NCI-H358 cells. For Spheroid formation assay, 1000 MIA PaCa-2 cells in tumorsphere media were seeded in each well of ultra-low attachment 6-well plates and treated the next day. For spheroid disintegration assay, 200 NCI-H358 cells in tumorsphere media were seeded in each well of ultra-low attachment plate and allowed to form spheroids before being subjected to drug treatment. Images are representative of three replicates.

The sensitivity of cells in 3D culture is believed to provide a more accurate prediction of *in vivo* efficacy, as it is highly correlated with drug response in xenograft models [28]. Therefore, we performed a spheroid formation assay, where treatment of MIA PaCa-2 cells in a 3D culture medium with a combination of KPT9274 with MRTX849 prevented the formation spheroids (**Figure 3B**). We further tested this combination in a spheroid disintegration assay, where KRAS G12C mutant NCI-H358 cells were first allowed to grow into sizeable spheroids in 3D culture medium and then exposed to the drugs for three days. The combination treatment resulted in enhanced disintegration of the spheroids (**Figure 3C**). These results demonstrate the efficacy of PAK4i and KRAS G12Ci combination in 3D cell growth models of KRASG12C mutant PDAC and NSCLC.

### Sotorasib and KPT9274 combination is more efficacious than single agent sotorasib in a KRASG12C mutant PDAC in vivo model

To assess the *in vivo* effect of sotorasib either as a single agent or in combination with KPT9274, a MIA PaCa-2 sub-cutaneous tumor xenograft model was established in ICR-SCID mice. The tumor-bearing mice were orally treated with sub-optimal doses of either KPT9274 (100 mg/Kg QD × 5 × 3 weeks), or sotorasib at one-fourth of MTD (25 mg/Kg QD x 5 × 3 weeks) or a combination of sotorasib (25 mg/Kg QD x 5 × 3 weeks) and KPT9274 (100 mg/Kg QD × 5 × 3 weeks). Oral administration of the combination led to significant reduction in tumor volumes (**Figure 4A**), tumor weights (**Figure 4B)** and tumor sizes (**Figure 4C)** as compared to single agent treatments. The drug treatments, either single agents or combination, caused no significant change in body weights of mice during the course of the treatment (**Figure 4D**). Also, we did not find any signs of either organ toxicity or metastatic spread while performing gross animal autopsy. Residual tumor profiling using IHC showed marked reduction in the proliferation marker Ki67, as well as inhibition of KRAS and PAK4 in the combination group (**Figure 4E**). In addition, Western Blot analysis of the residual tumor tissue proteins revealed downregulation of ERK activation as a result of treatment with KPT9274, or sotorasib and more so with their combination (**Figure 4F**). Expression of DUSP-6, a downstream target of ERK, was reduced in the tumor tissue from the combination group, as also the expression of cell cycle markers CDK4, CDK6 and cyclin D1, suggesting that the combination of PAK4i and KRASG12Ci induces cell cycle arrest at the G1/S phase in the tumor cells, ultimately resulting in tumor growth inhibition (**Figure 4G**). Additionally, the expression levels of PAK4 mRNA were found to be significantly decreased in the residual tumor from the combination group (**Figure 4H**). These results, taken together, demonstrate the preclinical safety and efficacy of sotorasib and KPT9274 combination *in vivo*.

**Figure 4.**
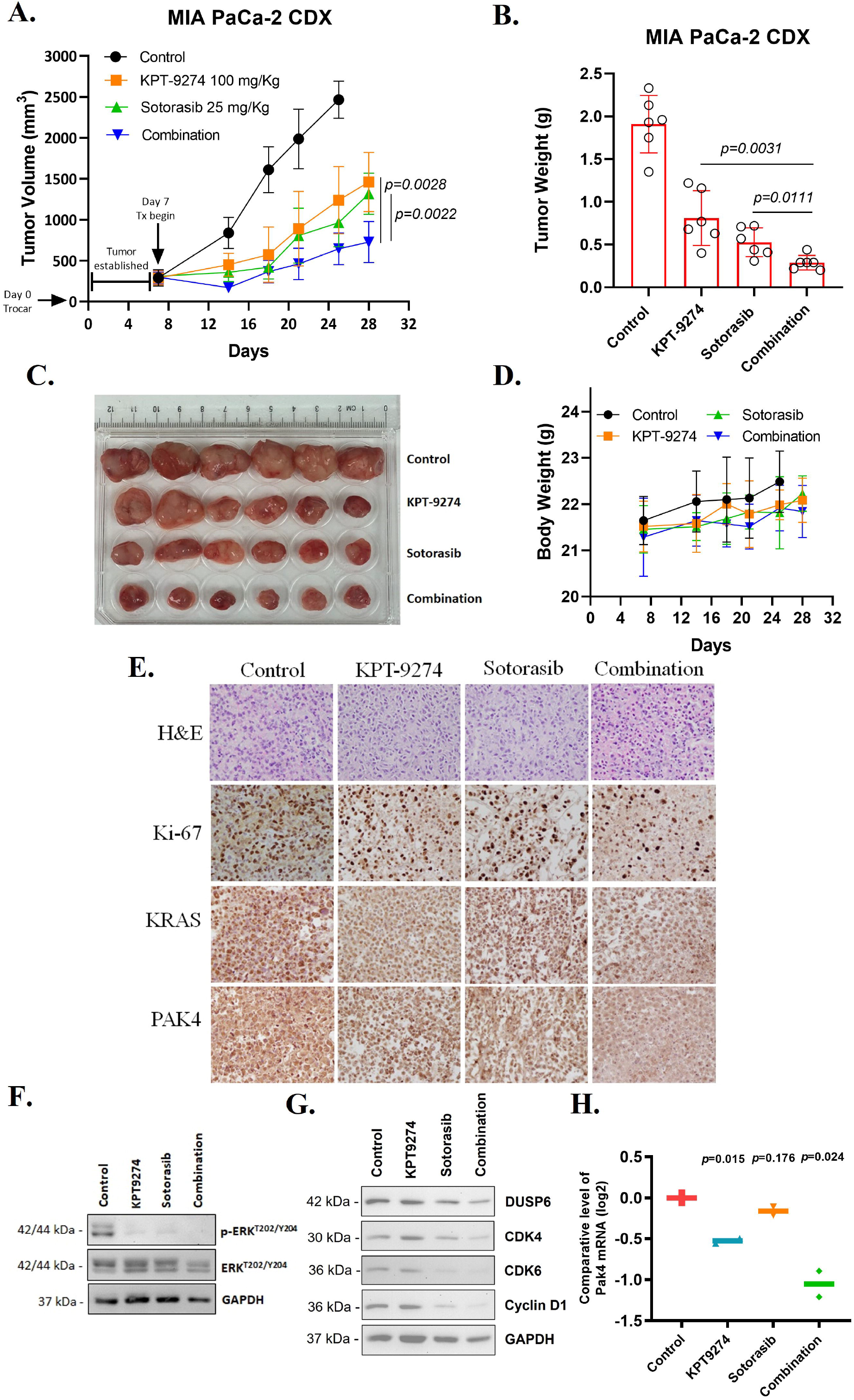
Preclinical antitumor efficacy of KPT9274 and KRASG12C inhibitor combination in a KRASG12C mutant PDAC CDX model. Treatment of ICR-SCID mice carrying subcutaneous KRASG12C mutant PDAC MIA PaCa-2 cell-derived tumor xenografts with a combination of KPT9274 and sotorasib resulted in significant reduction in (**A**) tumor volumes, (**B**) tumor weights, (**C**) and tumor sizes. A two-tailed unequal variance student’s *t* test was performed to statistically compare tumor volumes and tumor weights at the end of experiment. (**D**) No substantial loss in the animal body weights was observed. (**E**) IHC analysis showing (x400 magnification) reduction in Ki67, KRAS and PAK4 in tumor tissues harvested from mice treated with the combination of KPT9274 and sotorasib. (**F, G**) Immunoblots showing downregulation of p-ERK, its downstream target DUSP6, and cell cycle markers CDK4, CDK6 and cyclin D1 protein expression in the tumor tissue from the combination group. (**H**) The mRNA levels of PAK4 were also found decreased in the combination group as determined by RT-qPCR.

### KPT9274 is effective as an adjuvant to sotorasib for durable tumor growth inhibition and enhances survival in a KRASG12C mutant NSCLC in vivo model

In order to examine the effect of KPT9274 in preventing the relapse of tumors following the cessation of sotorasib treatment, we administered KPT9274, sotorasib, or their combination to mice harboring NCI-H358 tumor xenografts for 15 days, following which sotorasib treatment was withdrawn but treatment with KPT9274 was continued for an additional 11 days (**Figure 5A**). Tumors relapsed in mice treated with single agent sotorasib a few days after treatment was stopped. However, mice receiving KPT9274 in combination with, and later as an adjuvant to sotorasib therapy continued to show tumor remission. Significant reduction in tumor volumes, tumor weights and tumor sizes were observed in the combination compared to the single agent groups at the end of experiment (**Figure 5B-D**). The response in the combination/adjuvant arm of this *in vivo* study was durable and signifies not just the efficacy of KPT9274 in combination with KRASG12Ci but also highlights its utility as a maintenance therapy to prevent or delay tumor relapse.

**Figure 5.**
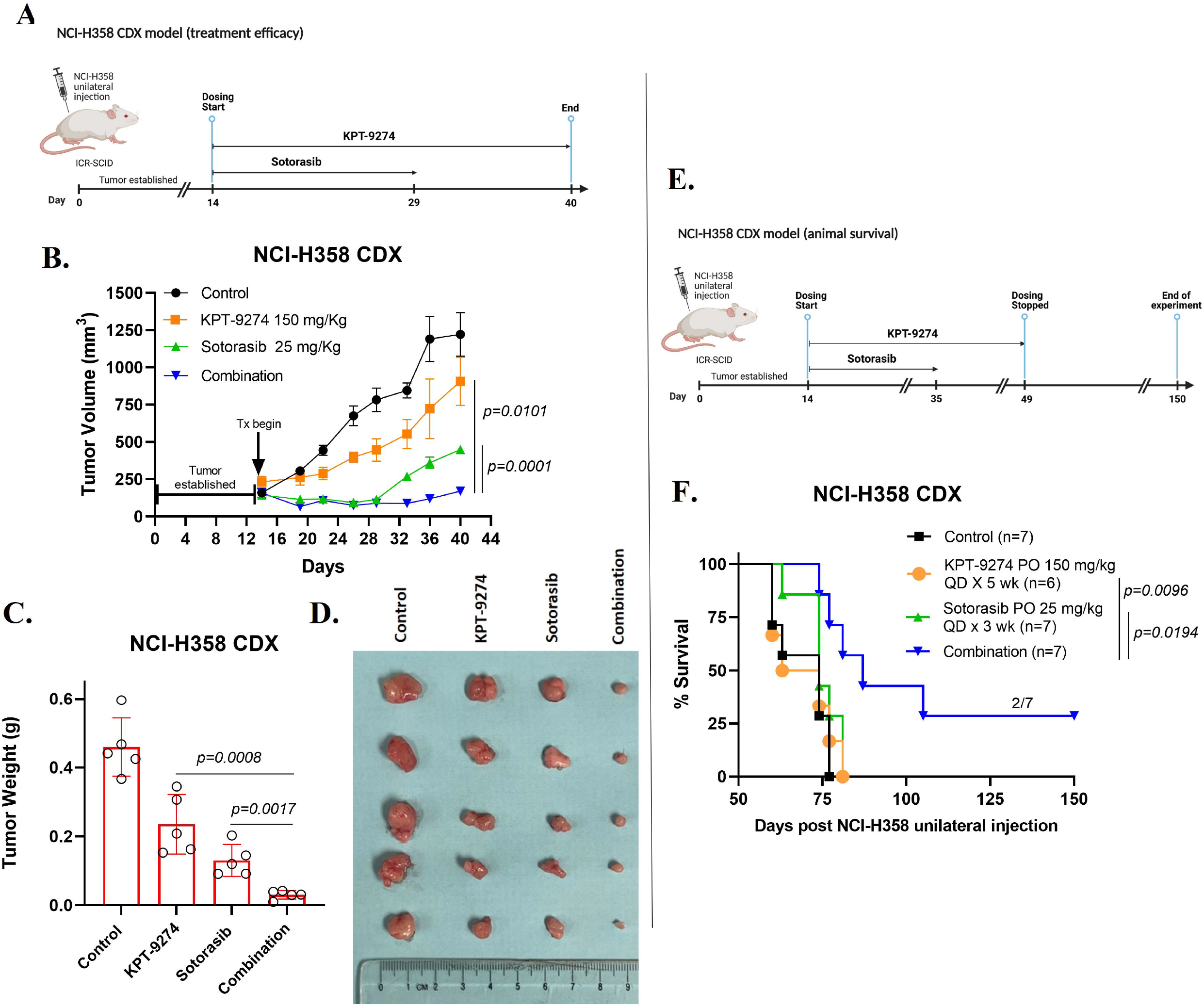
Combination/adjuvant therapy of KPT9274 with sotorasib induces durable tumor growth inhibition and enhances survival in a KRASG12C mutant NSCLC CDX model. (**A)** ICR-SCID mice subcutaneously engrafted with KRASG12C mutant NSCLC NCI-H358 cells were orally administered sotorasib (25 mg/Kg QD x 5) for 15 days in the single agent and combination groups, while KPT9274 (150 mg/Kg QD x 5) was dosed for 26 days in both the groups. This study timeline has been created with BioRender.com (License#*AC2561ROBN*). Treatment of mice carrying the CDX with KPT9274 and sotorasib combination results in significant decrease in (**B**) tumor volumes, (**C**) tumor weights, and (**D**) tumor sizes. A two-tailed unequal variance student’s *t* test was performed to statistically compare tumor volumes and tumor weights at the end of experiment. (**E)** In another experiment, mice unilaterally transplanted with NCI-H358 cells were administered with sotorasib (3 weeks), KPT9274 (5 weeks) or their combination daily five times a week by oral gavage. This study timeline has been created with BioRender.com (License#*UV2561UK0E*). (**F)** Two mice (29% survival) in the combination treatment group survived and remained tumor free till the culmination of the study i.e., 150 days post transplantation. Statistical comparison of survival curves was performed by Log-rank (Mantel-Cox) test.

KRASG12C mutant NSCLC NCI-H358 CDX model was again utilized to evaluate the effect of KPT9274 and sotorasib combination therapy on survival as an endpoint (**Figure 5E**). The results demonstrate a remarkable enhancement in survival of mice that received a combination of KPT9274 (150 mg/Kg QD x 5 × 5 weeks) and sotorasib (25 mg/Kg QD × 5 × 3 weeks). Almost 29% (2/7) of mice in the combination treatment group survived and remained tumor free for as long as 150 days post NCI-H358 subcutaneous transplantation (**Figure 5F**). This establishes that by incorporating KPT9274 as a combination/adjuvant partner to KRAS therapy, an increase in survival can be achieved.

## DISCUSSION

In this article, we show for the first time synergistic anticancer effects between KRASG12C inhibitors and a PAK4 inhibitor KPT9274. Our combination approach of co-targeting KRASG12C and PAK4 resulted in enhanced growth suppression of KRASG12C mutant cells and cell-derived xenografts (CDX). This study brings forward a novel combination therapy for drug resistant KRASG12C mutant tumors and provide preclinical rationale for the use of KPT9274 in a clinical setting to prevent or delay the development of resistance in patients receiving KRASG12C inhibitor monotherapy.

Targeting the KRAS oncoprotein has been a long-term objective in translational oncology. The development of inhibitors that can specifically target the KRASG12C protein has reignited hopes after decades of writing off KRAS as being non-targetable. Following the initial success of sotorasib and adagrasib in clinical trials [5,6], researchers in the field have doubled down on their efforts to target KRAS with renewed enthusiasm. The treatment options available to patients with the KRASG12C mutation have undoubtedly expanded with the FDA’s recent approval of sotorasib and adagrasib [7,8]. Despite showing promise as potent targeted therapies, KRASG12C inhibitors as monotherapies have limited efficacy. Studies have shown that patients treated with these inhibitors develop drug resistance over time, highlighting the need for combination approaches that could potentially enhance the sensitivity of tumors to KRAS inhibitors when co-targeted.

There is a growing understanding of the importance of identifying synthetic lethality associated with KRAS and developing small molecule inhibitors that target these synthetic lethal targets. Previously, we had experimentally validated synthetic lethal interactions of KRASG12C inhibitors with nuclear export protein XPO1 inhibitor *in silico, in vitro* and *in vivo* [26]. In this study, the identification of the existence of synthetic lethality between PAK4 and KRAS using the Slorth database rationalizes the significance of co-targeting PAK4 and KRAS.

PAK4 is activated through different signaling pathways in cancer and it also acts as a hub linking major oncogenic signaling pathways [9]. Approximately 20% of patients with pancreatic cancer exhibit gene amplification of PAK4, and PAK4 kinase activity is heightened in pancreatic tumors [29]. In NSCLC, overexpression of PAK4 is reportedly linked to metastasis, reduced survival rates, and an advanced stage of the disease [11]. Knockdown of PAK4 in KRAS mutant colon cancer cells suppresses their proliferation and this suppression was independent of Raf/MEK/ERK or PI3K/AKT signaling, indicating that some unidentified PAK4 effectors were involved [10]. While PAK4 mainly serves as a PI3K effector, it has also been reported to act as an upstream regulator of PI3K in driving resistance to cisplatin in gastric and cervical cancer cells [22,23], suggesting the existence of a feedback loop between Ras/PI3K and PAK4. Very recently, ablation of Kras oncogene has been shown to induce massive tumor regression and prevent the emergence of sotorasib resistance [30]. However, Kras ablated pancreatic cancer cells have earlier been reported to exhibit PI3K dependent MAPK signaling [31]. Being an effector of PI3K, PAK4 can serve as a potential pharmacological target, the inhibition of which can blunt the PI3K dependent survival signaling and thereby augment the efficacy of KRAS therapies. Based on this accumulating body of evidence, PAK4 has been implicated as an attractive target in cancers driven by mutant KRAS.

A combination therapy employing KRASG12C and PAK4 inhibitors can prove to be effective, especially considering that the KRASG12C inhibitor monotherapy has already been reported to result in the emergence of drug resistant subpopulations of cancer cells [32-34]. We hypothesized that using a PAK4 inhibitor in combination with a KRASG12C inhibitor would eliminate cancer cells that have developed resistance to the latter. This proposition was validated when we tested the PAK4 inhibitor KPT9274 against sotorasib-resistant cancer cell line (MIA-AMG-R) and found that MIA-AMG-R cells were indeed sensitive to KPT9274, even more than the parental KRASG12C mutant PDAC cells (MIA PaCa-2). Taken together, these results suggest that targeting PAK4 activity may serve as a viable therapeutic approach for overcoming resistance to KRASG12C inhibitors.

Inhibition of PAK4 by KPT9274 has been shown to enhance the efficacy of anti-PD-1 immunotherapy in a murine melanoma CDX model [35]. In a previous study, our group had demonstrated antitumor activity of KPT9274 in xenograft models of PDAC and its ability to overcome drug resistance and stemness [20]. In another study, pharmacological inhibition of PAK4 inhibited the growth of various human tumor cells both *in vitro* and *in vivo* [36]. The administration of PF3758309, another PAK4 inhibitor, significantly reduced the growth of colorectal tumors in patient-derived xenograft models of CRC [37]. Similarly, in this study we have shown that administering a PAK4 inhibitor in combination with a KRASG12C inhibitor results in extended survival and antitumor effects in KRASG12C mutant CDX models.

The results of our study indicate that combining a PAK4 inhibitor with a KRASG12C inhibitor can effectively suppress the growth of cancer cells carrying KRASG12C mutations in both 2D and 3D cell cultures. Moreover, this combination treatment substantially reduces the clonogenic potential of KRASG12C mutant cancer cells. Using KRASG12C mutant tumor xenograft models, we have demonstrated that the combination of KPT9274 and sotorasib is more effective at reducing tumor growth and improving survival rates. Our results also underscore the potential of KPT9274 to act as an adjuvant to sotorasib therapy. The underlying mechanisms for the antitumor effects involve the combination’s capacity to inhibit cell growth and survival signaling, and to impede cell cycle progression by suppressing CDK4/6 and cyclin D1 expression. The *in vivo* findings corroborate the *in vitro* results that KPT9274 treatment can sensitize cancer cells that have developed resistance to KRASG12C inhibitor. Further, these results also imply that combining KPT9274 with sotorasib (or other KRASG12C inhibitors) can have synergistic effects in cancer patients who have developed resistance to KRASG12C inhibitor therapy. To investigate this possibility, we have planned a Phase Ib/II clinical trial to evaluate the effectiveness of combining KPT9274 and sotorasib in patients who have not responded to sotorasib therapy. If successful, this innovative combination treatment could offer significant benefits for individuals with KRASG12C mutant cancers.

## Acknowledgements

Work in the Azmi lab is supported by NIH R37 grant R37CA215427 and NIH R01 grant R01CA24060701. The authors thank the SKY Foundation, and UCAN CER-VIVE Foundation for supporting part of this study.

