## Supplemental Figures for "Anticancer efficacy of KRASG12C inhibitors is potentiated by PAK4 inhibitor KPT9274 in preclinical models of KRASG12C mutant pancreatic and lung cancers"

### Slide 1
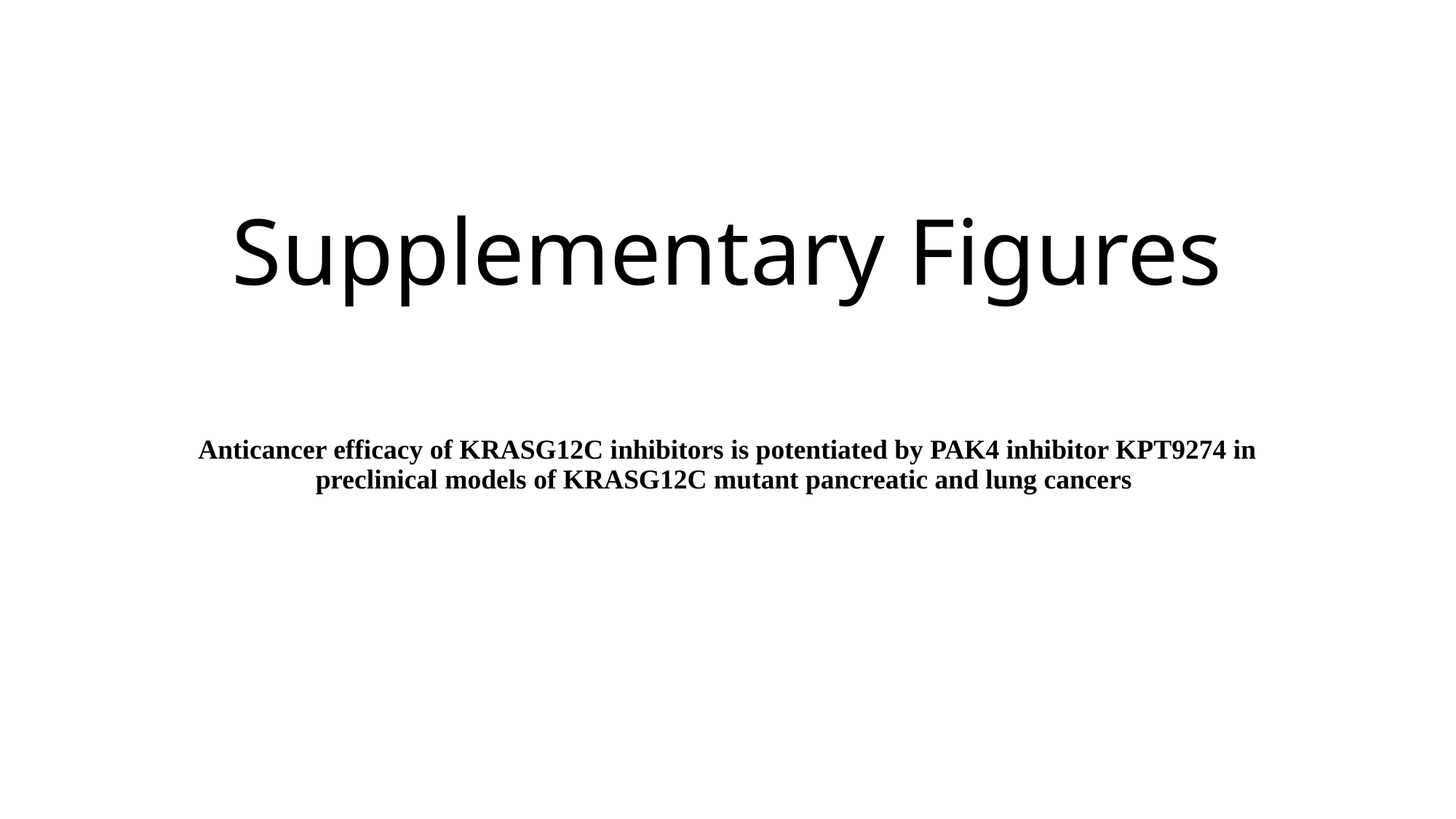

#### Supplementary Figures
Anticancer efficacy of KRASG12C inhibitors is potentiated by PAK4 inhibitor KPT9274 in preclinical models of KRASG12C mutant pancreatic and lung cancers

### Slide 2
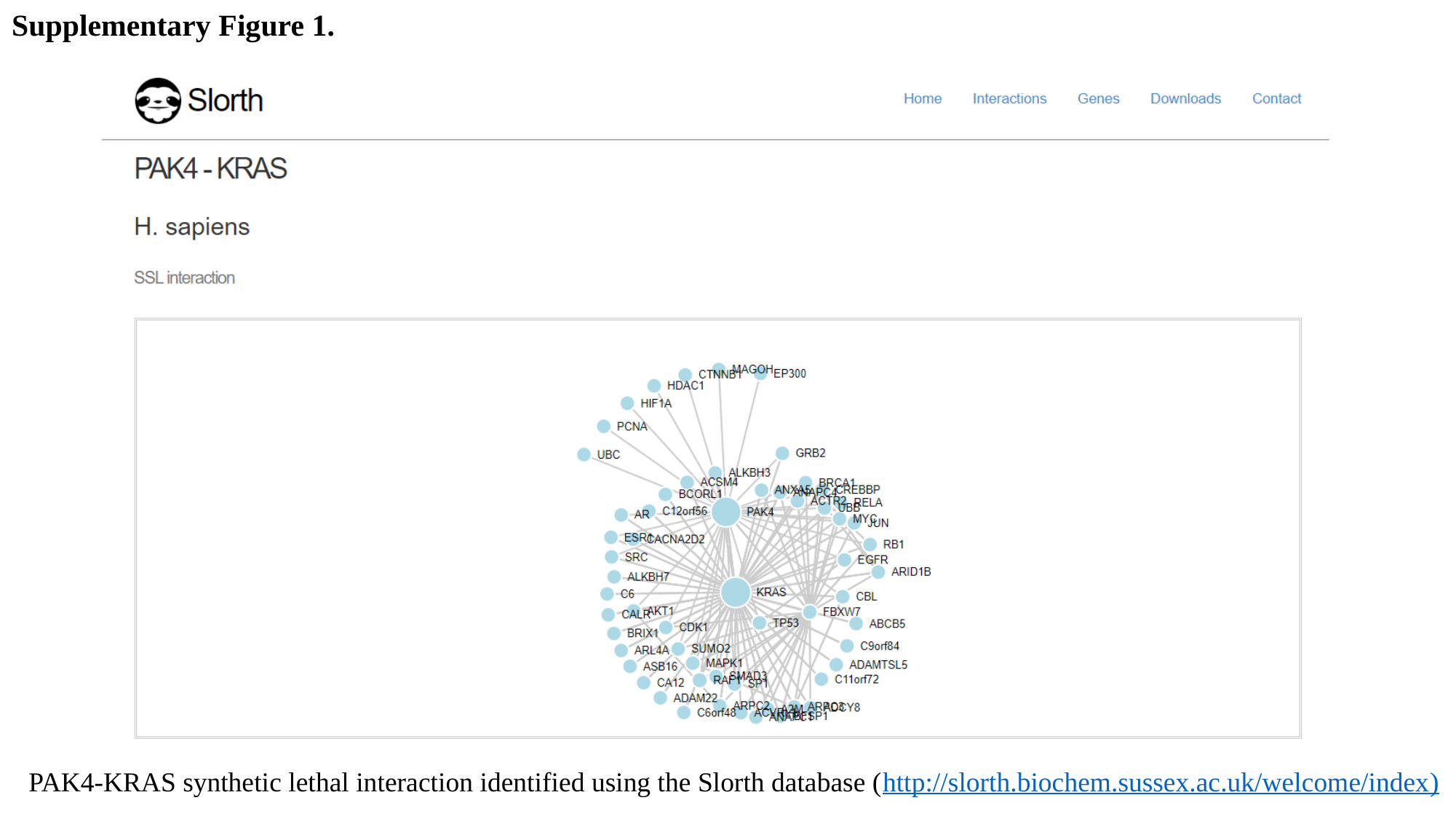

Supplementary Figure 1.
PAK4-KRAS synthetic lethal interaction identified using the Slorth database (http://slorth.biochem.sussex.ac.uk/welcome/index)

### Slide 3
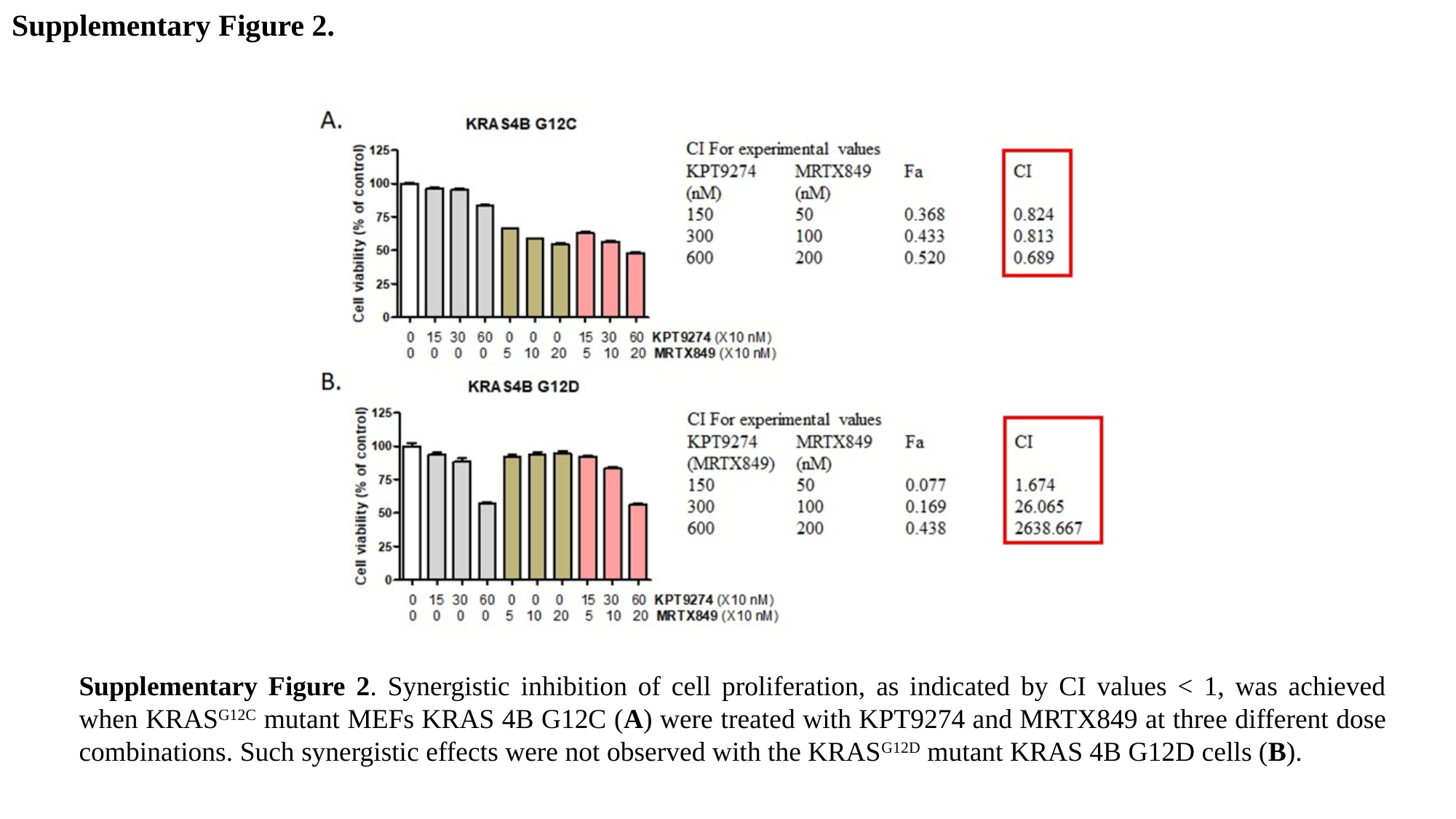

Supplementary Figure 2.
Supplementary Figure 2. Synergistic inhibition of cell proliferation, as indicated by CI values < 1, was achieved when KRAS^G12C^ mutant MEFs KRAS 4B G12C (A) were treated with KPT9274 and MRTX849 at three different dose combinations. Such synergistic effects were not observed with the KRAS^G12D^ mutant KRAS 4B G12D cells (B).
